# Changes in social reward across adolescence in male mice

**DOI:** 10.1101/2025.05.27.656363

**Authors:** Zofia Harda, Marta Klimczak, Klaudia Misiołek, Magdalena Chrószcz, Łukasz Szumiec, Maria Kaczmarczyk-Jarosz, Aleksandra Rzeszut, Rafał Ryguła, Barbara Ziółkowska, Jan Rodriguez Parkitna

## Abstract

In humans, adolescence is a time of dynamic behavioral and emotional changes, including a transient decrease in affect associated with being among family members. It is not clear if a similar change occurs in rodent species used to model human psychiatric disorders. Here, we investigated the developmental profile of the rewarding value of interactions with siblings across adolescence in male mice, using the social conditioned place preference task. We found that the reward value of social interactions followed a similar course to that in humans: high in early adolescence, it decreased in mid-adolescence and returned to the initial level in late adolescence. The observed change was specific to social interaction, as no age-dependent changes in preference for cocaine-conditioned context were detected. Taken together, these data show similarities between mice and humans in developmental changes in sensitivity to the rewarding effects of interactions with familiar kin.

## 1. Introduction

Adolescence is a time of rapid behavioral and neural changes, as well as the peak onset age for many mental disorders (Crone and Dahl, 2012; Kessler et al., 2005). It is postulated that the emergence of psychiatric symptoms during adolescence results from alterations in typical developmental processes (Paus et al., 2008). However, causal links between adolescent changes in brain maturation, behavior and pathophysiology have not been firmly established, partly because of the lack of proper animal models. It is thus of great importance to understand to what extent the behavioral development of model animals parallels the features of human adolescence (Lin and Wilbrecht, 2022).

In male laboratory mice, which descend from *Mus musculus* species, adolescence typically spans postnatal days (P) 30 to 60, corresponding to ages 11–21 years in humans (Bell, 2018). Although no precise biological markers define the start or end of adolescence in either species, significant physiological and neural changes occur during this period. For example, rapid increase in serum testosterone levels and nonlinear changes in the expression of D1 and D2 receptors in basal ganglia are observed (Clarkson et al., 2012; Cullity et al., 2019; Jean-Faucher et al., 1978). Despite these developmental dynamics, most behavioral studies treat adolescence as a uniform phase (Adriani et al., 1998; Moore et al., 2011; Shepard et al., 2017) or compare adults to a single adolescent timepoint, usually around P28-30 (Akers et al., 2012; Stone and Quartermain, 1997), with some notable exceptions (Nardou et al., 2019; Reiber et al., 2022). Here, we hypothesize that social reward in adolescent mice may also develop nonlinearly.

One of the characteristic features of human adolescence is changes in social preferences. Whereas infants fully depend on parental care and display strong distress following separation from their mothers, mid-adolescents show a decrease in the time spent with their family members along with an increase in the time spent alone or with peers(Larson and Richards, 1991; Nelson et al., 2005a). This behavioral shift is accompanied by changes in emotions associated with relatives: early and late adolescents show positive affect in the company of their family members, while mid-adolescents report more negative emotional states (Larson and Richards, 1991). However, it is not known whether affective changes resembling those observed in humans also occur in model animal species. Therefore, the goal of the present study was to assess the reward value of interactions with familiar kin in early, middle and late adolescence in male mice.

## 2. Methods and Materials

### 2.1. Animals

Experiments were performed with C57BL/6 male mice bred at the Maj Institute of Pharmacology animal facility. Mice were housed in a 12/12 h light-dark cycle (lights on at 7 AM) under controlled conditions: a temperature of 22 ± 2 °C and a humidity of 40-60%. After weaning, the mice were housed with all littermates of the same sex. Rodent chow and water were available ad libitum. Home and conditioning cages contained aspen nesting material and aspen gnawing blocks. Behavioral tests were conducted during the light phase under dim illumination (5-10 lux). Social conditioned place preference (sCPP) and social interaction tests were video recorded with additional infrared LED illumination. The age and weight of mice in each experimental group are summarized in **Table S1**.

Three age groups were studied: early (around postnatal day 33 [P33]), middle (P38) and late (P43) adolescence (for information about experimental groups, see **Tables S1** and **S2**).

All behavioral procedures were approved by the II Local Bioethics Committee in Krakow (permit numbers 35/2019, 185/2020, 38/2021, 266/2020, 305/2020) and performed in accordance with the Directive 2010/63/EU of the European Parliament and of the Council of 22 September 2010 on the protection of animals used for scientific purposes. The reporting in the manuscript follows the ARRIVE guidelines.

### 2.2 Social conditioned place preference

The test was performed to assess the rewarding effects of housing with siblings and followed the procedure described previously (Harda et al., 2022). The test consisted of three phases: pretest, conditioning, and posttest (**Fig. 1A**).

**Fig. 1.**
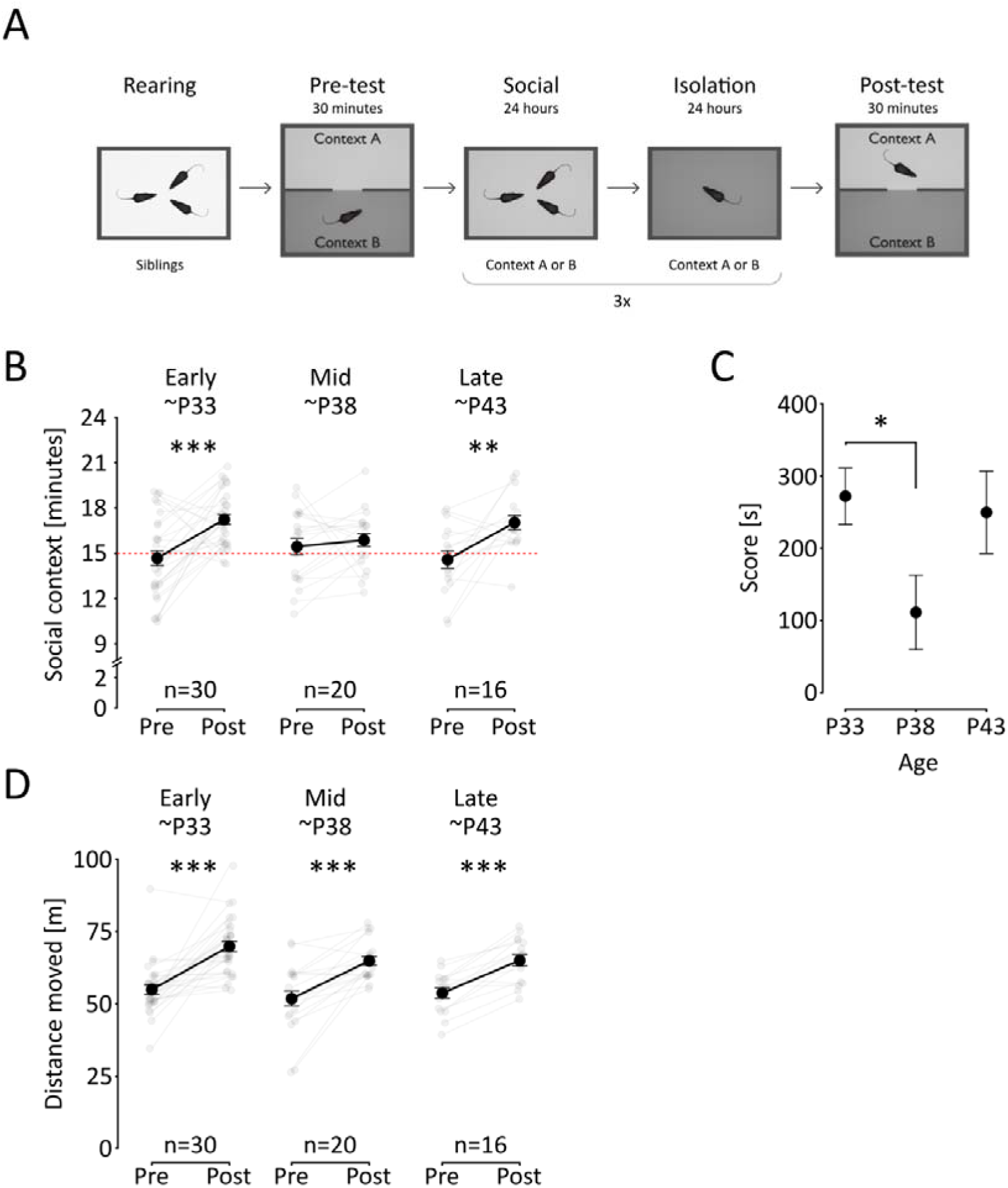
Social conditioned place preference during adolescence. (**A**) Schematic representation of the experimental schedule. **(B)** Time spent in the social context. The graphs show the mean time spent on the social-conditioned context during the pretest and posttest (black dots and line, error values are s.e.m.). The gray points and lines show all individual mice in the corresponding age bracket as indicated above. (C) Preference of the social context. The points show mean values of the preference score, i.e., the difference between the time spent in social context and isolation context during posttest of mice at different adolescence stages as indicated below. (D) Distance traveled during the tests. Points represent mean distance travelled during pretest and posttest, error bars are s.e.m., gray points and lines represent individual animals. Significant post-hoc (ANOVA with Šidák correction) differences between mean values are shown as: ‘**’ p <0.01, and ‘***’ p <0.001.

The pretest and posttest phases were performed in a two-compartment cage, as in previously published papers (Dölen et al., 2013; Harda et al., 2022; Hung et al., 2017; Misiołek et al., 2023; Nardou et al., 2019). Each cage compartment contained a novel context (context A or context B) defined by type of bedding and gnawing block size and shape. Bedding materials used were beech (context A, P.P.H. “WO-JAR”, Poland or PPHU Natur-Drew A. Czaja, Poland or Terrario Peak Wilderness, DMR Group, Poland) and cellulose (context B, Scott Pharma Solutions, cat no. L0107). In the home cages, aspen bedding was used (ABEDD, Latvia or Tapvei GLP, Estonia). Mice were allowed to freely explore the test cage for 30 minutes, and the time spent in each compartment was recorded. Animals that spent more than 70% of the pretest time in one of the contexts were excluded (**Table S2**).

After the pretest, animals were returned to their home cages for approximately 24 h. Then, mice were assigned to undergo social conditioning (housing with cage mates) for 24 h in one of the contexts used in the pretest followed by 24 h of isolate conditioning (single housing) in the other context. Conditioning was performed in cages identical to the home cage, with ad libitum access to food and water. To prevent bias, the social context was randomly assigned such that approximately half of the animals received social conditioning in context A and half in context B. In cases where the final number of animals conditioned in each context was not equal (due to an unequal number of animals passing the 70% criterion or unequal number of animals in the litter), we pseudorandomly trimmed the larger group using a Python script (https://zenodo.org/record/8100281)(Harda et al., 2025a). The exception from completely random selection was introduced to preserve a mean 50% initial context preference during the pretest, i.e., ascertain that the test was fully unbiased (**Table S4**). Analysis on the full data set (not trimmed) was also performed, to assess the potential interaction between the age and conditioning context on social reward (**Figure S1, Table S5**). The conditioning phase lasted 6 days (3 days in each context, alternating every 24 h), and then the posttest was performed.

Two measures of the rewarding effects of social interactions were used: 1) pretest vs. posttest comparison of the time spent in the social context, 2) score: time spent in the social context minus time spent in the isolation context during the posttest.

### 2.3. Social interaction in the partitioned cage

This test was carried out to assess social contact seeking with a sibling partner after 24 h of isolation. The procedure was performed in a rectangular cage (48 × 12 cm, 25 cm high) divided by a transparent, perforated plastic wall into two compartments: a smaller partner compartment and a larger focal animal compartment (**Fig. 2A**). One day before the test, the animals were weighed, and the heavier animal from each pair was designated as the focal animal. Next, the animals were habituated to their respective cage compartments for 10 minutes. During habituation, only one mouse was present in the test cage. After habituation, mice were placed in separate home cages for approximately 24 h, after which time the focal animals were placed in the test cage for the second adaptation session (5 minutes). After adaptation, the partner was introduced for 10 minutes. Two measures of social contact seeking were used: time spent in close proximity to the partner’s compartment and distance to the partner’s compartment.

**Fig. 2.**
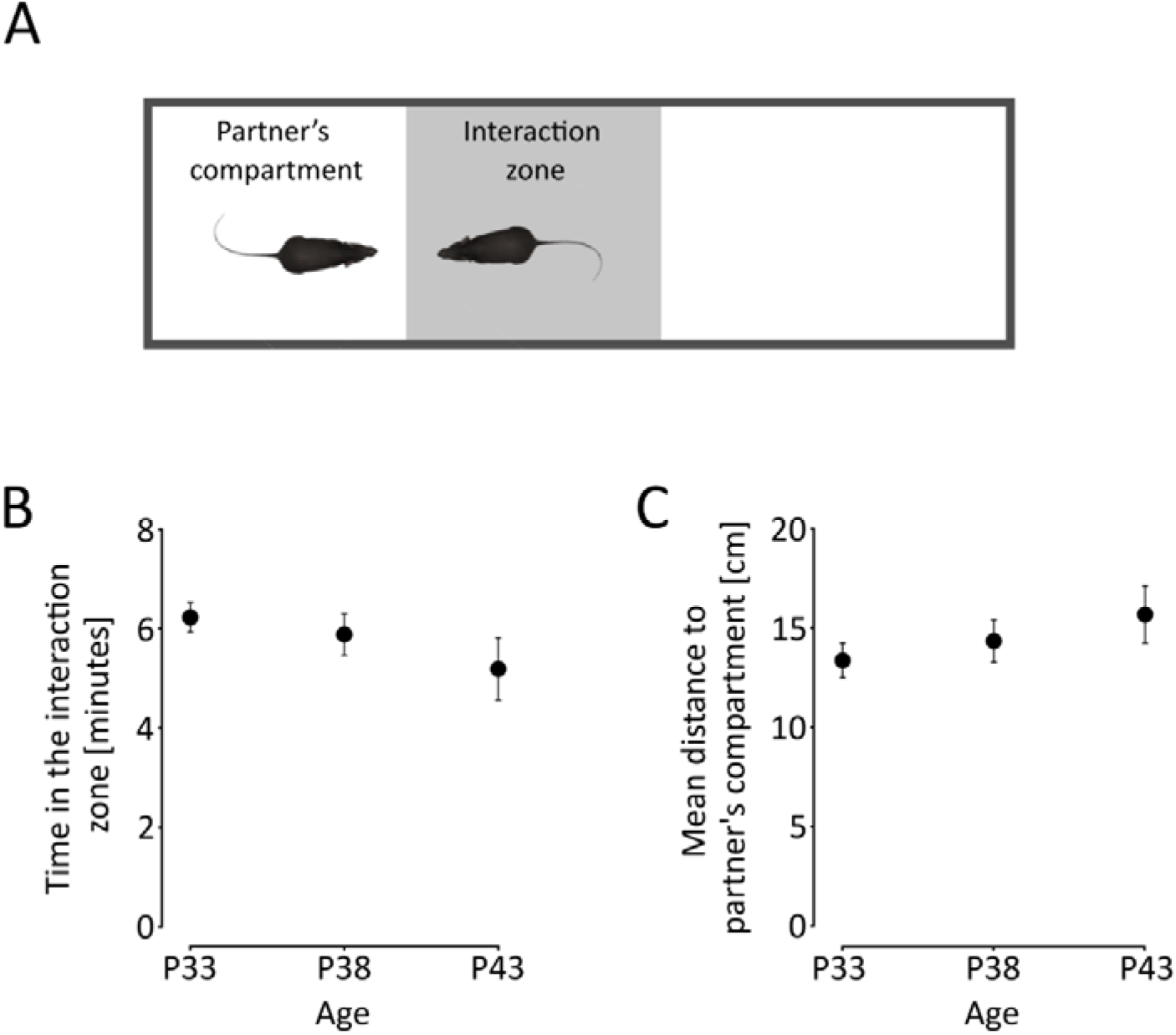
Social contact seeking during adolescence. (**A**) A schematic representation of the task. Mice were tested for seeking social contact with a sibling after 24 h of isolation. (**B**) Time spent in the interaction zone. The points show mean time spent in the zone, error bars are the s.e.m. (**C**) Mean distance from the partner’s compartment. Group sizes: P33 n = 10, P38 n = 18, P43 n = 10.

### 2.4. Cocaine-induced conditioned place preference

For the CPP paradigm, three-compartment cages were used (Med Associates, St. Albans, VT, USAMED-CPP-MSAT); the two peripheral compartments (that contained distinctive visual and tactile cues) were linked to the central compartment by guillotine doors. The test consisted of three phases: pretest, conditioning, and posttest. For the pretest and posttest phases, animals were introduced to the central compartment of the apparatus, and the doors between the compartments were lifted such that the animals could freely explore the apparatus for 20 minutes. Animals that spent more than 70% of the pretest time in one of the contexts were excluded. The less preferred of the two peripheral compartments was designated the cocaine compartment. The next day, the 3-day conditioning phase started. Each day, two 40-minute conditioning sessions were performed, separated by approximately 3 hours, during which animals were kept in their home cages. Before the morning conditioning session, animals received an i.p. saline injection, while before the afternoon session, they were injected with cocaine hydrochloride dissolved in saline (10 mg/kg, 5 μl/g). Immediately after the injection, animals were placed in the respective cage compartment, while the guillotine doors separating the compartments were closed. The posttest was performed on the day after the last conditioning session. The measures used to assess cocaine reward were the same as for the social conditioned place preference.

### 2.5. Data analysis

The distance traveled and time spent in separate cage compartments in the sCPP and social interaction tests were analyzed automatically using EthoVision XT 15 software (Noldus, The Netherlands). In the social interaction test, the zone close to the partner’s compartment was outlined digitally. In the cocaine-induced CPP test, the position of the mouse was registered automatically by the Med Associates system. Comparisons of sample means were performed using analysis of variance (ANOVA) followed by Sidak’s post hoc correction or Student’s t test for cases with only two samples. The statistical significance threshold was set at p < 0.05. Before the analysis, the Grubbs test for outliers was performed. For sCPP results, the outlier test was performed on “score”, “index 1”, “index 2”, and “distance moved during post-test” parameters before the trimming of the data. While only the “score” measure is shown in the present manuscript, all other measured are available at: https://zenodo.org/records/15284680.

The test was performed separately for the context A and B data. Four outliers were detected in the sCPP data: one in the P33 group, two in the P38 group, and one in the P44 group. For the social interaction data, the outlier test was performed on the parameter “time in interaction zone”. No outliers were detected. For the cocaine experiment, the outlier test was performed on “score”, “index 1”, “index 2”, “exploration during posttest”, and “movement during posttest” parameters. No outliers were detected. Data concerning animals excluded from the analysis are presented in **Table S2**. The analysis was performed in GraphPad Prism 9.4.1. For full descriptive statistics and statistical test results, see **Supplementary Material**.

## 3. Results

### 3.1. Rewarding effects of interactions with siblings across adolescence

To investigate the possible changes in the rewarding effects of social interactions with siblings during adolescence in mice, we used the social conditioned place preference (sCPP) test (**Fig. 1A)** (Harda et al., 2022) with male mice representing early (around postnatal day 33 [P33]), middle (P38) and late (P43) adolescence stages (for information about experimental groups, see **Tables S1** and **S2**). Animals were conditioned to associate one environmental context with group housing and another with social isolation, and then were tested to determine context preference. Mice aged 33, and 43 (but not 38) days at posttest showed a significant increase in the time spent in the social context from pretest to posttest, as revealed by significant interaction between age and conditioning (**Fig. 1B, Table S4,** Age F_2,63_ = 0.2182, p = 0.8046; Pre-Post F_1,63_ = 21.69, p < 0.0001; Age x Pre-Post F_2,63_ = 3.253, p = 0.0453; post hoc P33 p < 0.0001, P38 p = 0.8958, P43 p = 0.0064). There was no effect of context (i.e., the type of bedding) on preference (**Fig. S1, Table S5**). It had no significant effect as main factor or in interactions with age or conditioning, while interaction between age and conditioning remains significant when context is added as variable (Context F_1,65_ = 1.738, p = 0.1920; Age x Context F_2,65_ = 0.6916, p = 0.5044, Pre-Post x Context F_1,65_ = 0.1533, p = 0.6967; Age x Pre-Post x Context F_2,65_ = 0.2435, p = 0.7846; Age x Pre-Post F_2,65_ = 4.288 , p = 0.0178). Thus, interactions with siblings had lower reward value for mid-adolescent mice. The decrease in preference for the compartment associated with social contact in mid-adolescent mice was also clearly apparent in the social preference score (**Fig. 1C**). The score showed that the rewarding effects of social interactions were more than two times lower in mid-adolescent (P38) than in early-adolescent (P33) mice and returned to the early adolescent level in late-adolescent (P43) mice (Age F_2,63_ = 3.353, p = 0.0420). Importantly, motor activity was not significantly affected by age or interaction between age and conditioning; thus, it was not a confounding factor (**Fig. 1D, Table S6,** Age F_2,63_ = 2.279, p = 0.1108; Pre-Post F_1,63_ = 85.03, p < 0.0001; Age x Pre-Post F_2,63_ = 0.5909, p = 0.5569). Taken together, the results reveal a transient decrease in the rewarding effects of interactions with related individuals in mid-adolescent mice.

### 3.2. Social contact

Next, we investigated if the change in social behavior in mid-adolescent mice was specific to the rewarding effects of social interactions or if contact seeking was also altered. To explore this possibility, we administered a test in which contact with another mouse is enabled through a transparent, perforated plexiglass wall (**Fig. 2A**) (Langford et al., 2010). The interaction partners were siblings reared in the same cage but isolated for one day before the test to match the conditions of the sCPP posttest. We observed no age-related changes in the time spent in the proximity to the partner’s compartment (**Fig. 2B, Table S4,** Age F_2,65_ = 1.038, p = 0.3648) or the distance between the focal mouse and the partner’s compartment (**Fig. 2C**, Age F_1,65_ = 0.7805, p = 0.4660). These results might indicate that the decrease in the rewarding effects of interactions with related individuals in mid-adolescent mice was not accompanied by a general decrease in social contact seeking.

### 3.3. Rewarding effects of cocaine

An alternative explanation for the observed decrease in the rewarding effects of social interactions could be a general anhedonia or impairment in associative learning. To test whether the decrease in the rewarding effects of social interaction reflected a stimulus-independent reduction in the expression of conditioned behaviors, the cocaine-induced CPP was assessed. As expected, a significant increase in time spent in cocaine-paired compartment was observed but without the effect of age or interaction between age and conditioning (Fig. 3A, Table S4, Age F_2,41_ = 0.6877, p = 0.5084; Pre-Post F_1,41_ = 14.94, p = 0.004; Age x Pre-Post F_2,41_ = 0.3454, p = 0.7100). In line with this interpretation, no significant effect of age on the place preference score was detected (Fig. 3B, Age F_1,41_ = 0.9828, p = 0.3829). No significant effect of age or conditioning on locomotor activity was observed (Fig. 3C, Age F_2,41_ = 0.8940, p = 0.4168; Pre-Post F_1,41_ = 0.8799, p = 0.7682; Age x Pre-Post F_2,41_ = 0.8897, p = 0.4186). These data show that no general impairment of reward-conditioned preference occurred during mid-adolescence.

**Fig. 3.**
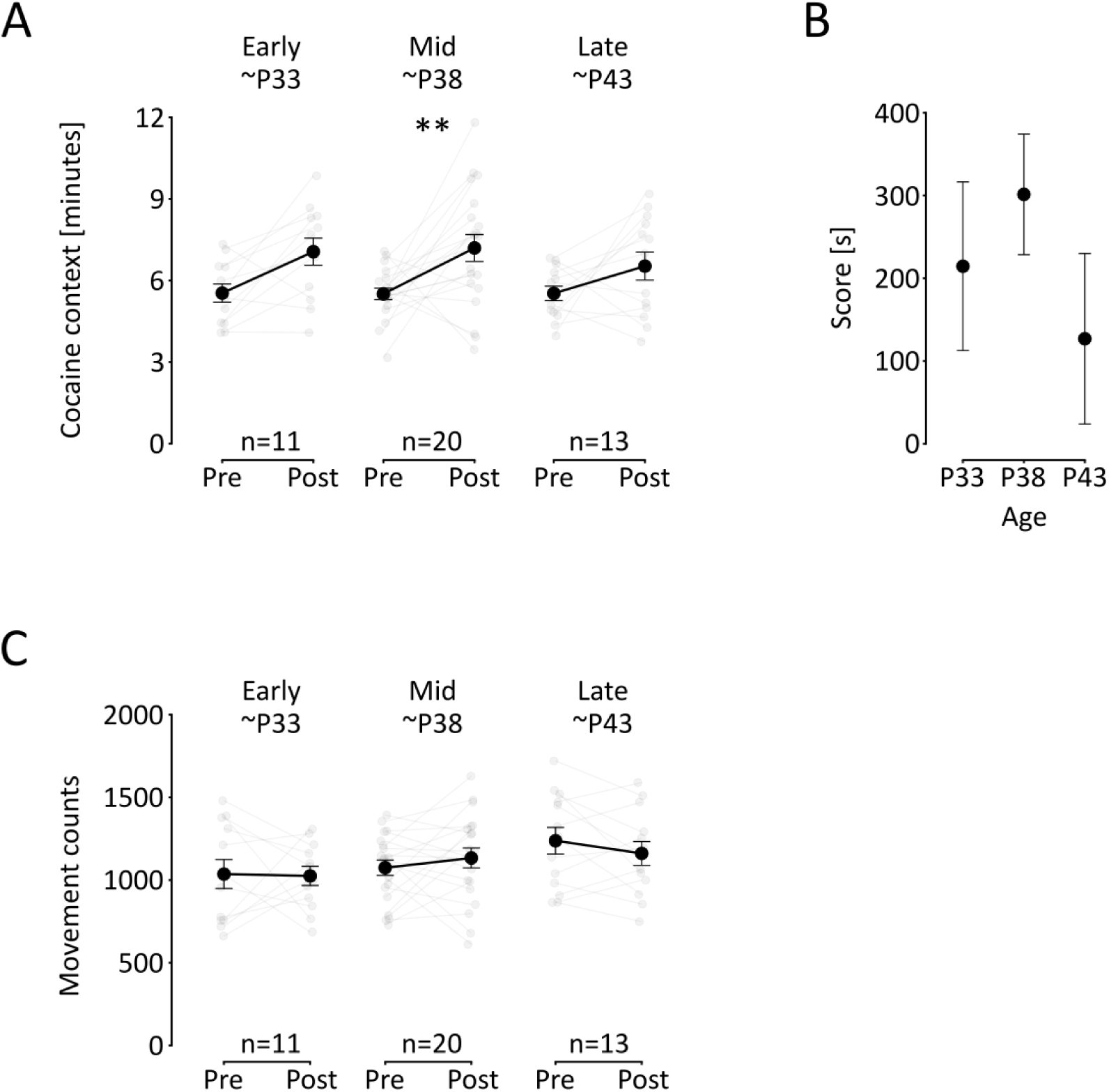
Cocaine-induced conditioned place preference during adolescence. (**A**) Time spent in the cocaine-conditioned context. The points show mean time spent in the cocaine-paired context during the pretest and posttest, error values are s.e.m.. The gray points and lines show individual mice in age brackets indicated above. (**B**) Preference of the cocaine-conditioned context. The points show mean values of the preference score, i.e. the difference between the time spent in cocaine-conditioned context and saline-paired context during posttest of animals at different adolescence stages as indicated below. (**C**) Distance traveled during the tests. Points represent mean distance travelled during pretest and posttest, error bars are s.e.m., gray points represent individual animals. A significant post-hoc (ANOVA with Šidák correction) difference between mean values is shown as: ‘**’ p <0.01.

## 4. Discussion

We found that the rewarding effects of interactions with familiar kin in male mice exhibit a transient decrease during mid-adolescence (around P36-40). This decrease was specific to social reward, as no change in social contact seeking, and the reward value of cocaine was observed during this period. Our results resemble human data reported by Larson and Richards (1991), who observed that the affect associated with time spent with family members was more positive in 10-year-old and 16-year-old boys than in 12- to 14-year-olds (Larson and Richards, 1991). The age range of 11 to 16 years in male humans corresponds to P30-40 in male mice, and this period is considered “peripubertal” (Bell, 2018). This may indicate that the phenomenon observed in our study, i.e., the temporary decrease in the reward value of interactions with familiar kin, is evolutionarily conserved. This finding may facilitate future research on the neuronal and physiological underpinnings of rebellious behaviors in adolescence.

Our results complement those of two previous studies that assessed the rewarding effects of social interactions using the sCPP paradigm at selected time points in adolescent mice (Cann et al., 2020; Nardou et al., 2019). In contrast to the methodology presented here, both previous studies used animals that were familiar but were not specifically kept in sibling groups. Our previous research on the reward value of social interactions has shown that eight weeks of familiarization with nonrelated mice is not equivalent to being reared in the same cage before weaning, at least in females, i.e. female mice do not form an sCPP when conditioned as adults in groups coming from different litters and housed together since weaning (Harda et al., 2022). Moreover, both previous studies (Cann et al., 2020; Nardou et al., 2019) used a paradigm with only two days of conditioning, which produces different results than the paradigm with six conditioning sessions used in this study (Harda et al., 2022; Misiołek et al., 2023). Bearing these differences in mind, we note that Cann and collaborators (2020) reported that social contact is rewarding in mice tested on postnatal day 29 but not on postnatal day 38, which aligns with our results. Conversely, in the study by Nardou and collaborators (2019), a decrease in the rewarding effects of social interactions during adolescence was not observed in male mice (although it was observed in females). However, the previous results were not interpreted in terms of the possible decrease in the reward value of social interactions during adolescence. Instead, Cann and collaborators conclude that sCPP, under the applied conditions, can be induced in early adolescent, but not adult mice, implying that the observed lack of conditioning effects at the age of 38 days is considered permanent. Results presented in this work, together with a previous paper where adult mice were studied (Misiołek et al., 2023), suggest that sCPP can be induced in late adolescent and adult male mice, provided that the 6-day protocol is used.

Recent study by Murray and collaborators (2024) investigated developmental changes in the motivation to obtain social reward in laboratory rats (Murray et al., 2024), a species in which the equivalent of P30-40 in mice is P42-55 (Bell, 2018). In this study, the operant conditioning protocol was employed and an unfamiliar conspecific was used as a stimulus animal. The authors found that social motivation is higher in pre-adolescent (P30) and mid-adolescent (P50) rats than in early adolescent animals (P40). This might indicate that the rewarding effects of and motivation to obtain social reward differ depending on the familiarity of the stimulus animal, with mid-adolescence being a period of decreased familiar kin contact seeking, and higher motivation to interact with unfamiliar individuals. A study using the 6-day sCPP protocol on mice housed with familiar and unfamiliar conspecifics in different adolescence timepoints should be performed to test this hypothesis. However, it must be noted that Murray and coworkers use different labels for the developmental periods, and describe P40 as mid-adolescence and P50 as late adolescence (Murray et al., 2024).

In contrast to social reward, there were no apparent changes in social contact seeking (assessed with the partition test) during adolescence. This finding indicates that the changes in social interactions in adolescent mice are qualitative rather than quantitative. We speculate that the amount of time spent with siblings may not change, but aggressive encounters may replace affiliative interactions. This interpretation is supported by earlier studies showing a profound decrease in passive social contact (Terranova et al., 1993) and play behavior (Wolff, 1981), along with an increase in fighting (Terranova et al., 1998, 1993), in the second postnatal month of mice life. Strikingly, the cocaine-induced CPP did not change during the adolescent period. These results indicate that the decrease in social CPP in mid-adolescent mice cannot be explained by a general decline in learning abilities and confirm that different neuronal processes underlie the rewarding effects of cocaine and social contact (Hung et al., 2017). Developmental changes in cocaine-induced CPP were previously studied in rats, and their results are inconsistent. For example, Brenhouse and coworkers reported a greater cocaine (10 mg/kg)-induced CPP in adolescents (P44) than in adults (P105) (Brenhouse et al., 2008). Notably, preadolescent rats (P27-37) showed similar levels of cocaine-induced CPP as adult animals (Brenhouse et al., 2008; Campbell et al., 2000), which suggests an inverted U-shaped relationship between cocaine reward and age. These results are consistent with previous reports of heightened sensitivity to other drugs of abuse in adolescent animals (Schramm-Sapyta et al., 2009). Conversely, the results by Badanich and collaborators indicated the highest sensitivity to cocaine reward in pre-adolescent rats, because a low cocaine dose (5 mg/kg) induced CPP only in pre-adolescents (P35), but not in adolescents (P45) or young adults (P60 animals) (Badanich et al., 2006). This suggests that some factors unidentified yet, for example the level of stress imposed by the experimental protocol, might differentially influence the rewarding effects of cocaine in different sub-periods of adolescence. Obviously, more studies are needed to resolve this issue. Irrespective of these inconsistencies, our results indicate that social and drug reward follow different developmental trajectories in the peri-adolescent period.

Although pubertal stage was not directly assessed in our study, previous research allows us to speculate on the physiological changes cooccurring with the observed behavioral shift. The time window analyzed in the present study (i.e. postnatal days 32-46) is a period of dramatic physiological changes associated with sexual development in mice. The first sign of sexual maturation in male mice—balano-preputial separation—appears between P28 and 32 (Manaserh et al., 2019; Zhou et al., 2012, n.d.). Plasma testosterone levels rise rapidly from P30 to P40 and begin to decline slightly after that age (Clarkson et al., 2012; Jean-Faucher et al., 1978). Neuroanatomical studies show rapid brain development during adolescence, including nonlinear changes in D1 and D2 dopamine receptor expression in the dorsal striatum, with the lowest D1/D2 ratio around PND 35 (Cullity et al., 2019). Physiological changes during puberty are accompanied with behavioral changes, especially in the social domain. Adolescence is a period of social re-orientation, characterized by the emergence of sexual behavior, decreased interest in family members, and increased interest in peers (Nelson et al., 2005b). Social re-orientation towards unfamiliar individuals is considered one of the causes of emigration from natal environment (dispersal). Dispersal begins around postnatal day 30 and continues well into adulthood, with approximately half of the males having migrated from their natal environment by postnatal day 42 (Groó et al., 2013). Our study suggests that a decrease in rewarding effects of interactions with familiar kin occurring in mid-adolescence might serve as a proximate affective mechanism of dispersal from the natal environment.

The main limitation of our study is that it was conducted exclusively on male mice. Female mice may exhibit a different developmental trajectory in the valuation of social rewards.

Taken together, our data show similarities between male mice and humans in the pattern of social reward development.

## Author contributions

Conceptualization: ZH, BZ, RR, JRP; Methodology: ZH, KM, MK, MC and JRP; Investigation: ZH, MK, KM, MC, ŁS, AR and MKJ; Visualization: ZH, KM and JRP; Supervision: BZ, JRP; Writing—original draft: ZH and BZ; Writing—review & editing: ZH, BZ, RR, and JRP with help from all the authors.

## Data availability

All data are available at https://zenodo.org/records/15284680. This paper is available at bioRxiv (Harda et al., 2025b).

## Declaration of generative AI and AI-assisted technologies in the writing process

During the preparation of this work the authors used ChatGPT in order to correct grammar mistakes and shorten two paragraphs. After using this tool, the authors reviewed and edited the content as needed and take full responsibility for the content of the published article.

## Supplementary Material

### Supplementary Results

Social Conditioned Place Preference

As a result of trimming procedure (see section 2. Methods) no deviation from the chance value (i.e. 50%) in the time spent in social context during the pre-test was detected in any of the groups, which confirms that the test was fully unbiased (Table S6).

**Fig. S1.**
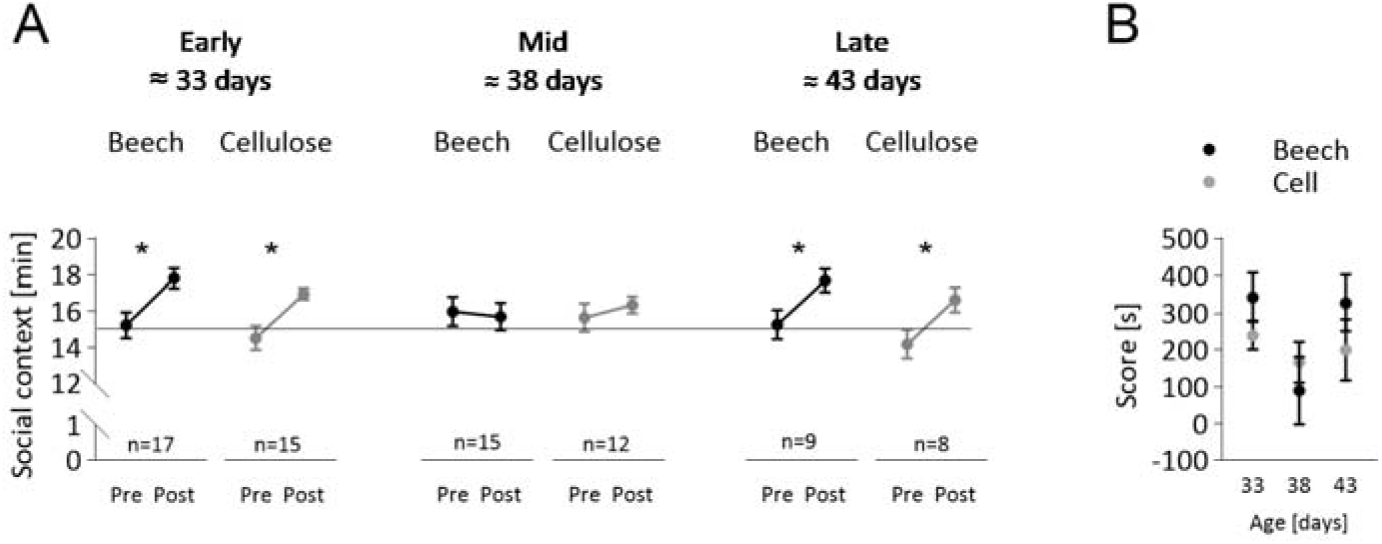
Effects of bedding type in sCPP protocol. (**A**) No effect of conditioning context type (or interaction between the context and age) on the time spent in the social context was detected. (**B**) No effect of conditioning context type on social preference score was detected. This figure uses the same dataset that was used on Figure 1 in the Main Text. ‘*’ p <0.05, ‘**’ p <0.01, and ‘***’ p <0.001.

**Table S1.** Age and weight of animals used in the study.

| Behavioral test | Figure | Age group [days] | n | Age [days] |  |  | Weight pre-test [g] |  |  | Weight post-test [g] |  |  |
| --- | --- | --- | --- | --- | --- | --- | --- | --- | --- | --- | --- | --- |
|  |  |  |  | Range | Mean | SEM | Range | Mean | SEM | Range | Mean | SEM |
| sCPP | 1 | ≈33 | 30 | 32-35 | 33.97 | 0.2222 | 7.3-16 | 10.38 | 0.3913 | 12-17.3 | 14.9 | 0.3733 |
|  |  | ≈38 | 20 | 38-39 | 38.05 | 0.5000 | 10.7-17.7 | 14.68 | 0.4411 | 15.7-21.1 | 18.22 | 0.3368 |
|  |  | ≈43 | 16 | 43-44 | 43.63 | 0.1250 | 12.4-21 | 17.4 | 0.6862 | 15.9-21.1 | 18.69 | 0.4408 |
| Social interaction in partition cage | 2 | 33 | 10 | 33-33 | 33 | 0 | NA | NA | NA | 11.6-18.8 | 16.09 | 0.7289 |
|  |  | ≈38 | 18 | 36-39 | 37.5 | 0.3638 | NA | NA | NA | 13.5-22 | 17.85 | 0.6241 |
|  |  | 42 | 10 | 42-42 | 42 | 0 | NA | NA | NA | 16.6-22 | 19.67 | 0.598 |
| Cocaine CPP | 3 | ≈33 | 12 | 32-33 | 32.17 | 0.1124 | 8.5-13.1 | 11.34 | 0.3963 | 10.8-16.4 | 14.13 | 0.4916 |
|  |  | ≈38 | 20 | 36-40 | 37.6 | 0.2938 | 13.4-20.5 | 16.96 | 0.5141 | 17-21.2 | 19.07 | 0.2865 |
|  |  | ≈43 | 13 | 41-46 | 43.23 | 0.4551 | 16-21.8 | 19.42 | 0.4397 | 16.4-22.1 | 19.39 | 0.4594 |

**Table S2.** Animals excluded from the analysis.

| Behavioral test | Figure | n initial | n excluded<br>based on<br>initial<br>context<br>preference<br>(> 70%) | n died | n<br>excluded<br>due to<br>health<br>reasons | n<br>excluded<br>due to<br>technical<br>mistakes | n<br>excluded<br>based<br>on<br>outlier<br>test | n<br>excluded<br>by<br>Python<br>script to<br>equalize<br>the<br>number<br>of<br>animals<br>on the<br>two<br>social<br>contexts | n<br>analyzed |
| --- | --- | --- | --- | --- | --- | --- | --- | --- | --- |
| sCPP | 1 | 78 | 2 | 0 | 1 | 0 | 4 | 5 | 66 |
| Social interaction in partition cage | 2 | 38 | NA | 0 | 0 | 0 | 0 | NA | 38 |
| Cocaine CPP | 3 | 47 | 2 | 0 | 0 | 1 | 0 | NA | 44 |

**Table S3.** Social conditioned place preference: descriptive statistics and one-sample t test results (refers to Main Text Figure 1).

**Main Text Figure 1).**
| Age<br>group<br>[days] | df | Pre-test time spent in social<br>context [minutes] |  |  |  | Post-test time spent in social<br>context [minutes] |  |  |  | Score [s] |  |  |  |
| --- | --- | --- | --- | --- | --- | --- | --- | --- | --- | --- | --- | --- | --- |
|  |  | Mean | SEM | One-sample t<br>test (difference<br>from chance<br>value) |  | Mean | SEM | One-sample t<br>test (difference<br>from chance<br>value) |  | Mean | SEM | One-sample t<br>test (difference<br>from chance<br>value) |  |
|  |  |  |  | t | p-value |  |  | t | p-value |  |  | t | p-value |
| ≈33 | 29 | 14.67 | 0.4882 | 0.6721 | 0.5068 | 17.23 | 0.3272 | 6.817 | <0.0001 | 272 | 39 | 6.924 | <0.0001 |
| ≈38 | 19 | 15.43 | 0.5353 | 0.8093 | 0.4283 | 15.87 | 0.4273 | 2.027 | 0.057 | 111 | 52 | 2.157 | 0.044 |
| ≈43 | 15 | 14.57 | 0.5799 | 0.7369 | 0.4725 | 17.02 | 0.4740 | 4.271 | 0.0007 | 250 | 57 | 4.385 | 0.0005 |

**Table S4.**
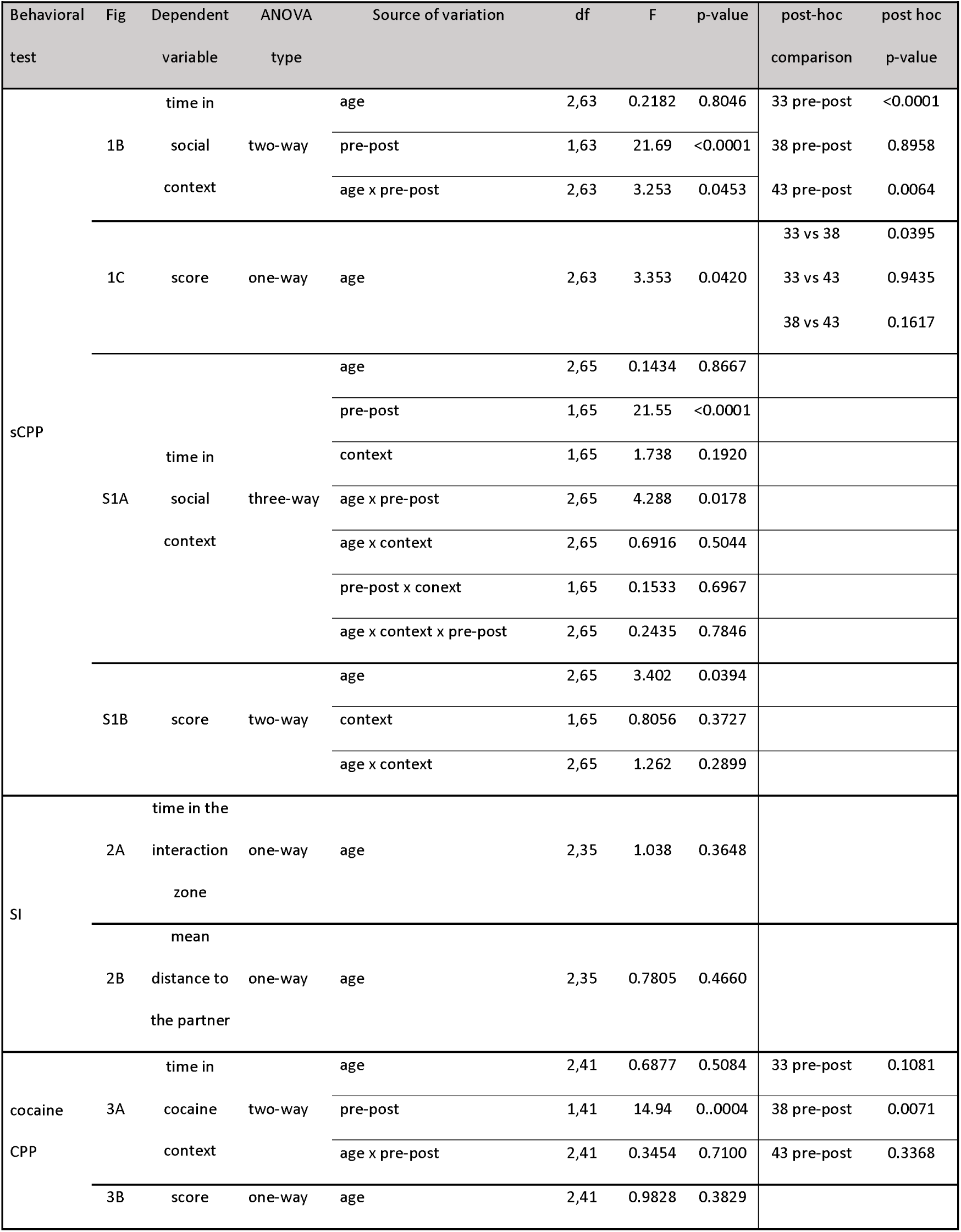
sCPP, social interaction and cocaine CPP tests: ANOVA results.

**Table S5.** Effects of the conditioning context (bedding type) on the sCPP results in different age groups: descriptive statistics and one-sample t-test (refers to Figure S1 in the Supplementary Information)

**descriptive statistics and one-sample t-test (refers to Figure S1 in the Supplementary Information)**
| Social context | Age group<br>[days] | df | Pre-test time spent in social context |  |  |  | Post-test time spent in social context [minutes] |  |  |  | Score [s] |  |  |  |
| --- | --- | --- | --- | --- | --- | --- | --- | --- | --- | --- | --- | --- | --- | --- |
|  |  |  | [minutes] |  |  |  |  |  |  |  |  |  |  |  |
|  |  |  | Mean | SEM | One-sample t test<br>(difference from<br>chance value) |  | Mean | SEM | One-sample t test<br>(difference from<br>chance value) |  | Mean | SEM | One-sample t<br>test (difference<br>from chance<br>value) |  |
|  |  |  |  |  | t | p-value |  |  | t | p-value |  |  | t | p-value |
| Beech | ≈33 | 16 | 15.24 | 0.7146 | 0.3318 | 0.7443 | 17.82 | 0.5528 | 5.108 | 0.0001 | 343 | 66.28 | 5.179 | <0.0001 |
|  | ≈38 | 9 | 15.98 | 0.8045 | 1.212 | 0.2563 | 15.69 | 0.7552 | 0.9182 | 0.3825 | 90 | 91.65 | 0.9862 | 0.3498 |
|  | ≈43 | 8 | 15.24 | 0.8091 | 0.3002 | 0.7717 | 17.68 | 0.6359 | 4.212 | 0.0029 | 328 | 76.40 | 4.294 | 0.0026 |
| Cellu-<br>lose | ≈33 | 14 | 14.53 | 0.6629 | 0.7129 | 0.4876 | 16.97 | 0.3259 | 6.047 | <0.0001 | 241 | 39.16 | 6.140 | <0.0001 |
|  | ≈38 | 11 | 15.63 | 0.7829 | 0.8025 | 0.4393 | 16.34 | 0.4642 | 2.882 | 0.0149 | 167 | 55.09 | 3.035 | 0.0114 |
|  | ≈43 | 7 | 14.17 | 0.7983 | 1.051 | 0.3282 | 16.62 | 0.6907 | 2.346 | 0.0514 | 201 | 82.76 | 2.427 | 0.0456 |

**Table S6.**
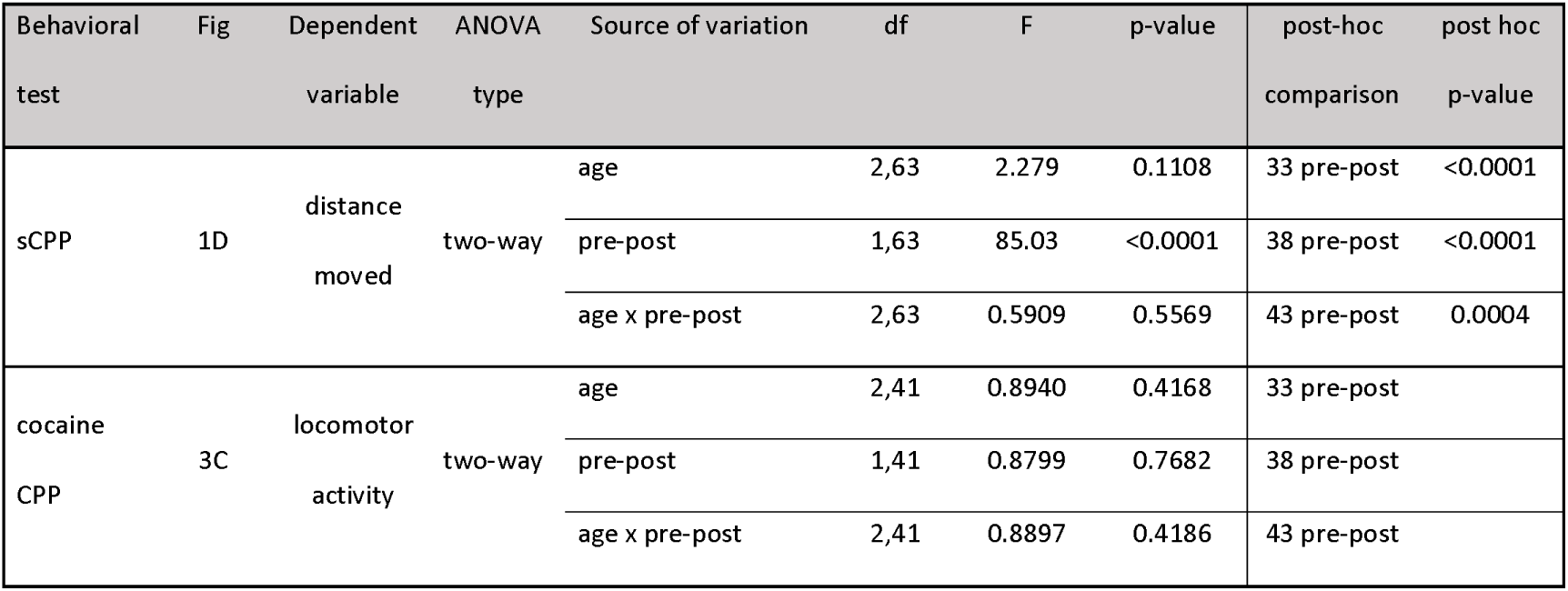
Locomotor activity during the sCPP and cocaine CPP tests: ANOVA (refers to Figure 1 and 3 the Main Text).

## References

Adriani, W., Chiarotti, F., Laviola, G., 1998. Elevated novelty seeking and peculiar d-amphetamine sensitization in periadolescent mice compared with adult mice. Behavioral Neuroscience 112, 1152–1166. 10.1037/0735-7044.112.5.1152

Akers, K.G., Arruda-Carvalho, M., Josselyn, S.A., Frankland, P.W., 2012. Ontogeny of contextual fear memory formation, specificity, and persistence in mice. Learn. Mem. 19, 598–604. 10.1101/lm.027581.112

Badanich, K.A., Adler, K.J., Kirstein, C.L., 2006. Adolescents differ from adults in cocaine conditioned place preference and cocaine-induced dopamine in the nucleus accumbens septi. Eur J Pharmacol 550, 95–106. 10.1016/j.ejphar.2006.08.034

Bell, M.R., 2018. Comparing Postnatal Development of Gonadal Hormones and Associated Social Behaviors in Rats, Mice, and Humans. Endocrinology 159, 2596–2613. 10.1210/en.2018-00220

Brenhouse, H.C., Sonntag, K.C., Andersen, S.L., 2008. Transient D1 Dopamine Receptor Expression on Prefrontal Cortex Projection Neurons: Relationship to Enhanced Motivational Salience of Drug Cues in Adolescence. J Neurosci 28, 2375–2382. 10.1523/JNEUROSCI.5064-07.2008

Campbell, J.O., Wood, R.D., Spear, L.P., 2000. Cocaine and morphine-induced place conditioning in adolescent and adult rats. Physiology & Behavior 68, 487–493. 10.1016/S0031-9384(99)00225-5

Cann, C., Venniro, M., Hope, B.T., Ramsey, L.A., 2020. Parametric investigation of social place preference in adolescent mice. Behav Neurosci 134, 435–443. 10.1037/bne0000406

Clarkson, J., Shamas, S., Mallinson, S., Herbison, A.E., 2012. Gonadal Steroid Induction of Kisspeptin Peptide Expression in the Rostral Periventricular Area of the Third Ventricle During Postnatal Development in the Male Mouse. Journal of Neuroendocrinology 24, 907–915. 10.1111/j.1365-2826.2012.02294.x

Crone, E.A., Dahl, R.E., 2012. Understanding adolescence as a period of social–affective engagement and goal flexibility. Nature Reviews Neuroscience 13, 636–650. 10.1038/nrn3313

Cullity, E.R., Madsen, H.B., Perry, C.J., Kim, J.H., 2019. Postnatal developmental trajectory of dopamine receptor 1 and 2 expression in cortical and striatal brain regions. J of Comparative Neurology 527, 1039–1055. 10.1002/cne.24574

Dölen, G., Darvishzadeh, A., Huang, K.W., Malenka, R.C., 2013. Social reward requires coordinated activity of accumbens oxytocin and 5HT. Nature 501, 179–184. 10.1038/nature12518

Groó, Z., Szenczi, P., Bánszegi, O., Altbäcker, V., 2013. Natal dispersal in two mice species with contrasting social systems. Behav Ecol Sociobiol 67, 235–242. 10.1007/s00265-012-1443-z

Harda, Z., Chrószcz, M., Misiołek, K., Klimczak, M., Szumiec, Ł., Kaczmarczyk-Jarosz, M., Rodriguez Parkitna, J., 2022. Establishment of a social conditioned place preference paradigm for the study of social reward in female mice. Sci Rep 12, 11271. 10.1038/s41598-022-15427-9

Harda, Z., Klimczak, M., Misiołek, K., Chrószcz, M., Rzeszut, A., Szumiec, Ł., Kaczmarczyk-Jarosz, M., Ryguła, R., Ziółkowska, B., Parkitna, J.R., 2025a. Mu and delta opioid receptor antagonists increase the expression of social conditioned place preference in early adolescent mice. 10.1101/2023.07.19.549691

Harda, Z., Klimczak, M., Misiołek, K., Chrószcz, M., Szumiec, Ł., Kaczmarczyk-Jarosz, M., Rzeszut, A., Ryguła, R., Ziółkowska, B., Parkitna, J.R., 2025b. Changes in social reward across adolescence in male mice. 10.1101/2025.05.27.656363

Hung, L.W., Neuner, S., Polepalli, J.S., Beier, K.T., Wright, M., Walsh, J.J., Lewis, E.M., Luo, L., Deisseroth, K., Dölen, G., Malenka, R.C., 2017. Gating of social reward by oxytocin in the ventral tegmental area. Science 357, 1406–1411. 10.1126/science.aan4994

Jean-Faucher, C., Berger, M., de Turckheim, M., Veyssiere, G., Jean, C., 1978. Developmental patterns of plasma and testicular testosterone in mice from birth to adulthood. Acta Endocrinol (Copenh) 89, 780–788. 10.1530/acta.0.0890780

Kessler, R.C., Berglund, P., Demler, O., Jin, R., Merikangas, K.R., Walters, E.E., 2005. Lifetime prevalence and age-of-onset distributions of DSM-IV disorders in the National Comorbidity Survey Replication. Arch Gen Psychiatry 62, 593–602. 10.1001/archpsyc.62.6.593

Langford, D.J., Tuttle, A.H., Brown, K., Deschenes, S., Fischer, D.B., Mutso, A., Root, K.C., Sotocinal, S.G., Stern, M.A., Mogil, J.S., Sternberg, W.F., 2010. Social approach to pain in laboratory mice. Soc Neurosci 5, 163–170. 10.1080/17470910903216609

Larson, R., Richards, M.H., 1991. Daily companionship in late childhood and early adolescence: changing developmental contexts. Child Dev 62, 284–300.

Lin, W.C., Wilbrecht, L., 2022. Making sense of strengths and weaknesses observed in adolescent laboratory rodents. Current Opinion in Psychology 45. 10.1016/j.copsyc.2021.12.009

Manaserh, I.H., Chikkamenahalli, L., Ravi, S., Dube, P.R., Park, J.J., Hill, J.W., 2019. Ablating astrocyte insulin receptors leads to delayed puberty and hypogonadism in mice. PLOS Biology 17, e3000189. 10.1371/journal.pbio.3000189

Misiołek, K., Klimczak, M., Chrószcz, M., Szumiec, Ł., Bryksa, A., Przyborowicz, K., Rodriguez Parkitna, J., Harda, Z., 2023. Prosocial behavior, social reward and affective state discrimination in adult male and female mice. Sci Rep 13, 5583. 10.1038/s41598-023-32682-6

Moore, E.M., Linsenbardt, D.N., Melón, L.C., Boehm II, S.L., 2011. Ontogenetic differences in adolescent and adult C57BL/6J and DBA/2J mice: Anxiety-like, locomotor, and consummatory behaviors. Developmental Psychobiology 53, 141–156. 10.1002/dev.20501

Murray, S.H., Logan, R.J., Sheehan, A.C., Paolone, A.R., McCormick, C.M., 2024. Developmental trajectory of social reward motivation from early adolescence into adulthood in female and male Long-Evans rats. Dev Psychobiol 66, e22495. 10.1002/dev.22495

Nardou, R., Lewis, E.M., Rothhaas, R., Xu, R., Yang, A., Boyden, E., Dölen, G., 2019. Oxytocin-dependent reopening of a social reward learning critical period with MDMA. Nature 569, 116–120. 10.1038/s41586-019-1075-9

Nelson, E.E., Leibenluft, E., McClure, E.B., Pine, D.S., 2005a. The social re-orientation of adolescence: a neuroscience perspective on the process and its relation to psychopathology. Psychol Med 35, 163–174. 10.1017/s0033291704003915

Nelson, E.E., Leibenluft, E., McClure, E.B., Pine, D.S., 2005b. The social re-orientation of adolescence: a neuroscience perspective on the process and its relation to psychopathology. Psychol Med 35, 163–174. 10.1017/s0033291704003915

Paus, T., Keshavan, M., Giedd, J.N., 2008. Why do many psychiatric disorders emerge during adolescence? Nat Rev Neurosci 9, 947–957. 10.1038/nrn2513

Reiber, M., Koska, I., Pace, C., Schönhoff, K., von Schumann, L., Palme, R., Potschka, H., 2022. Development of behavioral patterns in young C57BL/6J mice: a home cage-based study. Sci Rep 12, 2550. 10.1038/s41598-022-06395-1

Schramm-Sapyta, N.L., Walker, Q.D., Caster, J.M., Levin, E.D., Kuhn, C.M., 2009. Are adolescents more vulnerable to drug addiction than adults? Evidence from animal models. Psychopharmacology (Berl) 206, 1–21. 10.1007/s00213-009-1585-5

Shepard, R., Beckett, E., Coutellier, L., 2017. Assessment of the acquisition of executive function during the transition from adolescence to adulthood in male and female mice. Developmental Cognitive Neuroscience 28, 29–40. 10.1016/j.dcn.2017.10.009

Stone, E.A., Quartermain, D., 1997. Greater Behavioral Effects of Stress in Immature as Compared to Mature Male Mice. Physiology & Behavior 63, 143–145. 10.1016/S0031-9384(97)00366-1

Terranova, M.L., Laviola, G., Alleva, E., 1993. Ontogeny of amicable social behavior in the mouse: Gender differences and ongoing isolation outcomes. Developmental Psychobiology 26, 467–481. 10.1002/dev.420260805

Terranova, M.L., Laviola, G., de Acetis, L., Alleva, E., 1998. A description of the ontogeny of mouse agonistic behavior. J Comp Psychol 112, 3–12. 10.1037/0735-7036.112.1.3

Wolff, R., 1981. Solitary and social play in wild Mus musculus (Mammalia). Journal of Zoology 195, 405–412.

Zhou, Y., Guan, Q., Li, K., Tao, L., Hu, J., Xiao, J., 2012. Dissection of the maternal effects on puberty onset by embryo transplantation in mouse. J Endocrinol Invest 35, 676–680. 10.3275/8125

Zhou, Y., Zhu, W., Guo, Z., Zhao, Y., Song, Z., Xiao, J., n.d. Effects of maternal nuclear genome on the timing of puberty in mice offspring. Journal of Endocrinology 193, 405–412. 10.1677/joe.1.07049

